## Supplementary Materials for "Implications of the three-dimensional chromatin organization for genome evolution in a fungal plant pathogen"

### SUPPLEMENTARY DATA

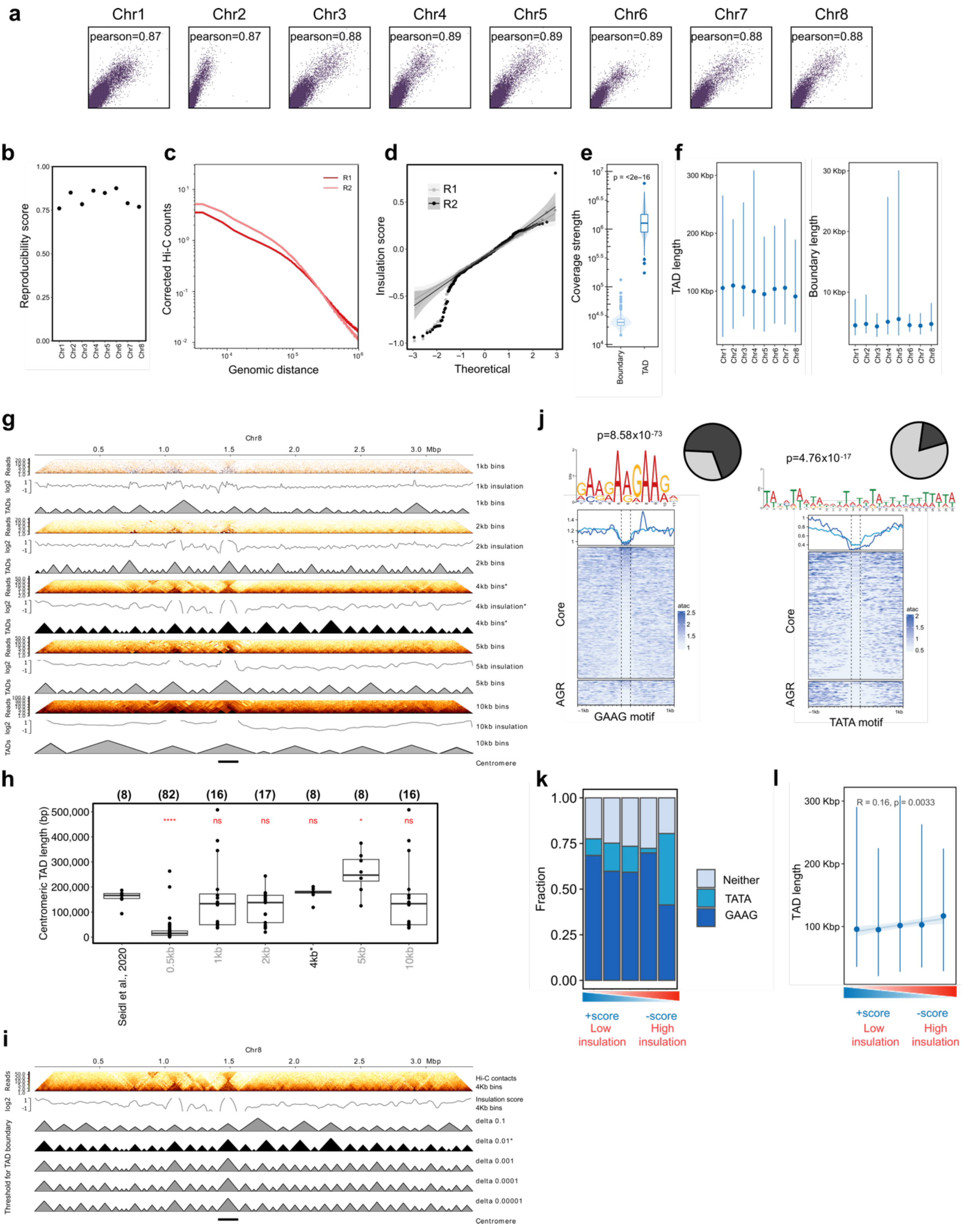


**Figure S1. Characteristics of TADs and TAD boundaries in *Verticillium dahliae* strain JR2. (a)** Pearson correlation between replicates of Hi-C interactions for *V. dahliae* strain JR2 divided per chromosome. **(b)** Reproducibility score based on a stratified cross-correlation of interactions between replicates of Hi-C interactions for *V. dahliae* strain JR2 divided per chromosome. **(c)** Distance-dependent interaction frequency in replicates of Hi-C data for *V. dahliae* strain JR2. **(d)** Quantile-Quantile plot of insulation scores calculated for each Hi-C replicate, confirming non-significant differences between replicates. **(e)** Coverage strength (Hi-C counts) over boundaries and bodies of TADs in merged replicates, showing that boundaries are depleted of interactions. P-value after Wilcoxon rank sum test. **(f)** Length of TADs bodies and boundaries predicted for the merged replicates, divided per chromosome. **(g)** Hi-C interactions normalized at different bins for Chromosome 8. From top to bottom for each bin strategy: Hi-C interaction matrix, insulation score and TAD prediction. In black the 4kb bin strategy used for further analyses. (**h**) Centromeric TADs prediction at different binning strategies, compared to centromeric regions defined by CenH3 and repeat content by Seidl et al., 2020. P-value after one-way Wilcoxon rank sum test. Upper numbers in brackets depict the total number of centromeric TADs predicted at each binning strategy. (**i**) TAD prediction at different threshold for bin insulation values using the Hi-C interaction matrix normalized at 4kb for Chromosome 8. In black the TAD prediction at the delta used for further analyses. (**j**) A GAAG-motif (left) and TATA-motif (right) is enriched in TAD boundaries of *V. dahliae* strain JR2 and correlate with a decrease in chromatin accessibility as determined by the assay for transposase accessible chromatin (ATAC). The top plot displays the average ATAC signal over TATA motifs with 1 kb up- and down-stream sequence in the core genome (light blue) and in AGRs (dark blue). Heatmap display ATAC signal around the GAAG and TATA motifs as rows in the core genome and in AGRs, respectively. **(k)** Occurrence of the GAAG-motif and the TATA-motif in boundaries separated based on their insulation score. **(l)** Correlation between boundary insulation score and length of adjacent TAD for boundary quintiles separated based on their insulation score.

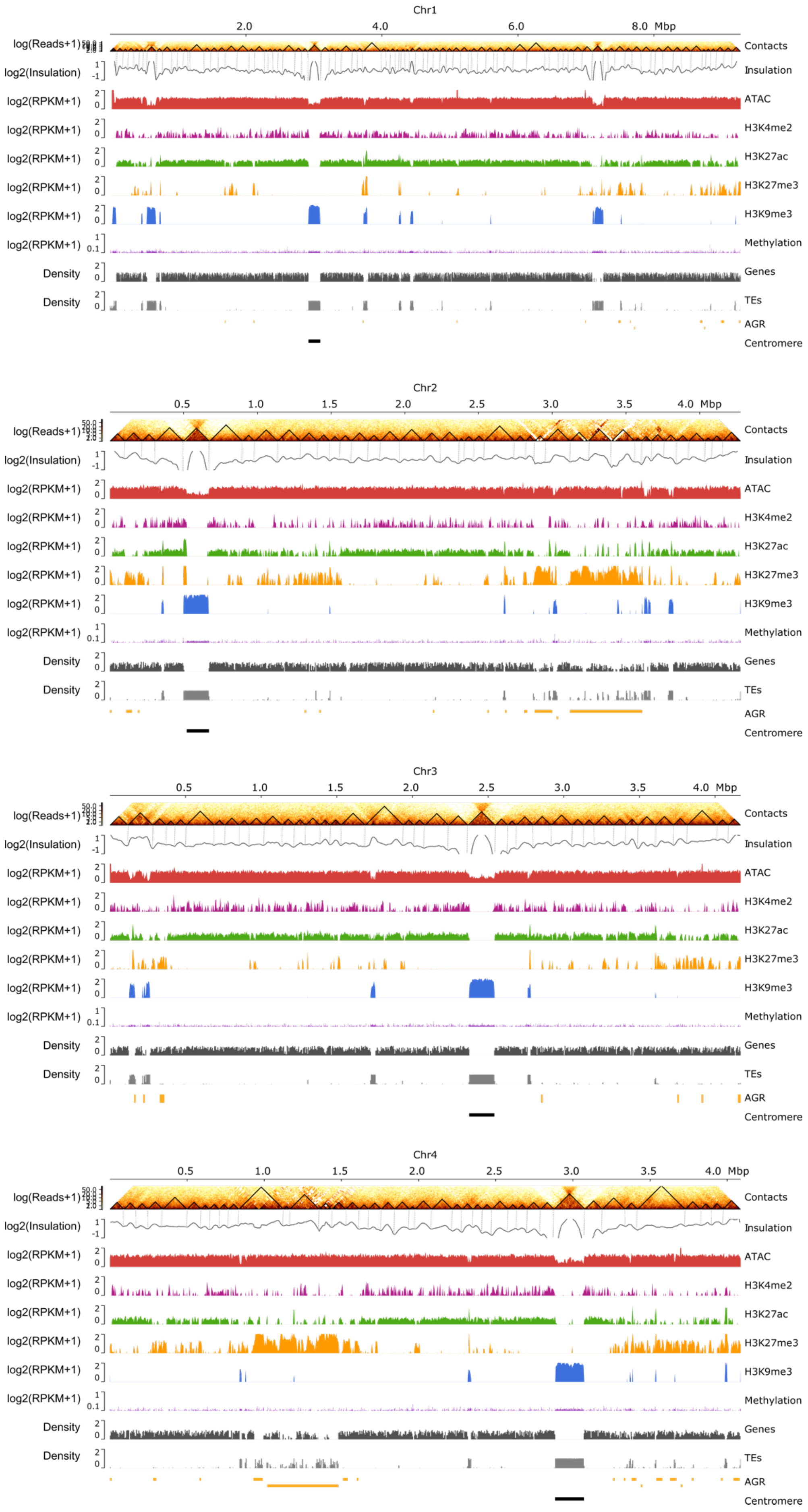

Chr1
2.0
4.0
6.0
8.0 Mbp
log(Reads+1)
Contacts
log2(Insulation)
Insulation
log2(RPKM+1)
ATAC
H3K4me2
H3K27ac
H3K27me3
H3K9me3
Methylation
Density
Genes
TEs
AGR
Centromere
Chr2
0.5
1.0
1.5
2.0
2.5
3.0
3.5
4.0 Mbp
Chr3
Chr4

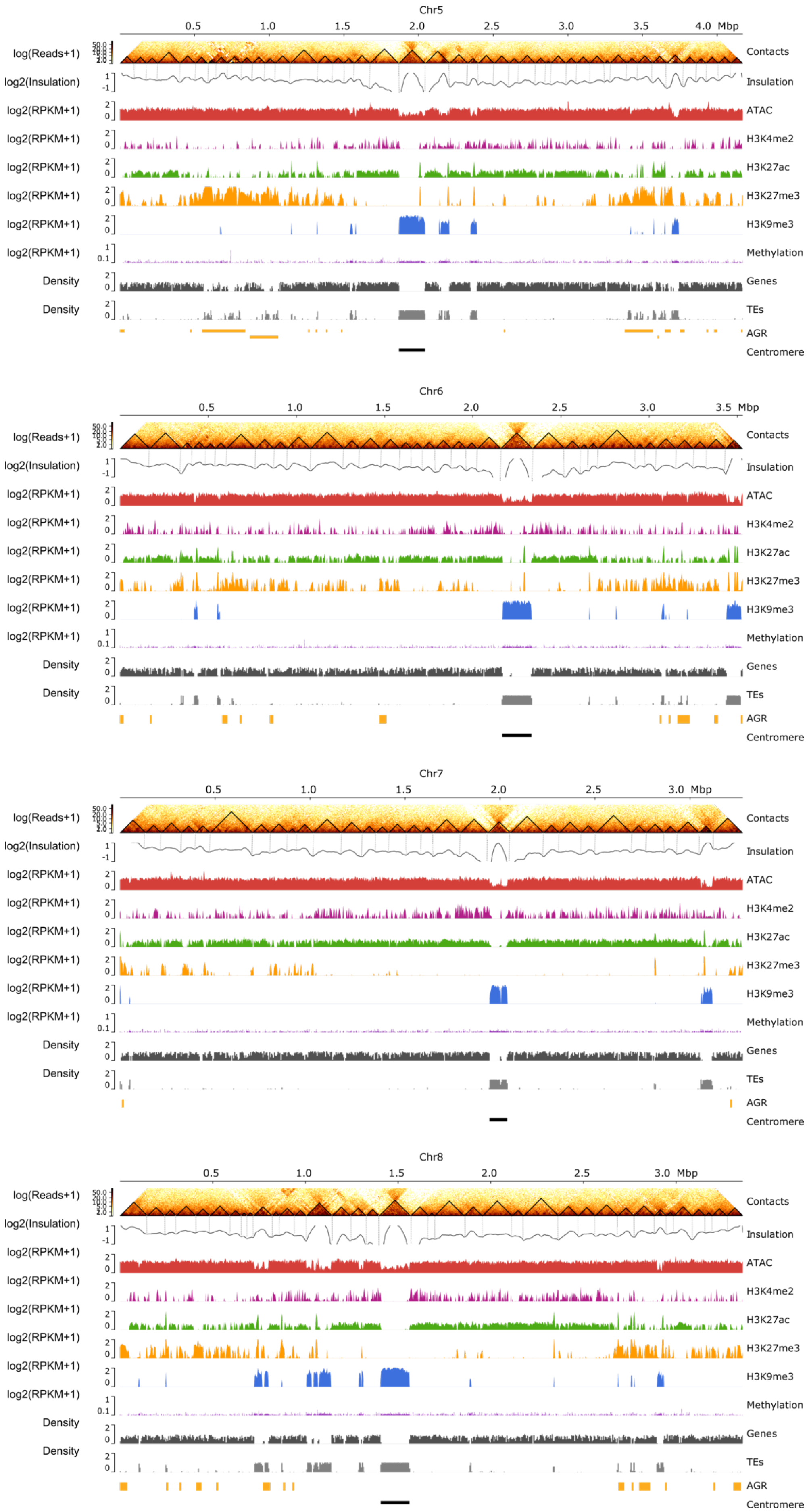
Chr5
0.5
1.0
1.5
2.0
2.5
3.0
3.5
4.0 Mbp
log(Reads+1)
log2(Insulation)
log2(RPKM+1)
Density
Contacts
Insulation
ATAC
H3K4me2
H3K27ac
H3K27me3
H3K9me3
Methylation
Genes
TEs
AGR
Centromere
Chr6
3.5 Mbp
Chr7
3.0 Mbp
Chr8
3.0 Mbp

**Figure S2. Distribution of TADs over all chromosomes of *Verticillium dahliae* strain JR2.** From top to bottom: Chromosome sizes, Hi-C contact matrix depicting TADs as black triangles, insulation score mapping and vertical dashed lines showing the insulation decrease associated to TAD boundaries, open chromatin regions as determined with ATAC-seq, histone modifications H3K4me2, H3K27ac, H3K27me3, and H3K9me3 normalized over a micrococcal nuclease digestion control, DNA methylation, gene and transposable element (TE) densities in 10 kb windows, and adaptive genomic regions (AGRs) and centromeric regions.

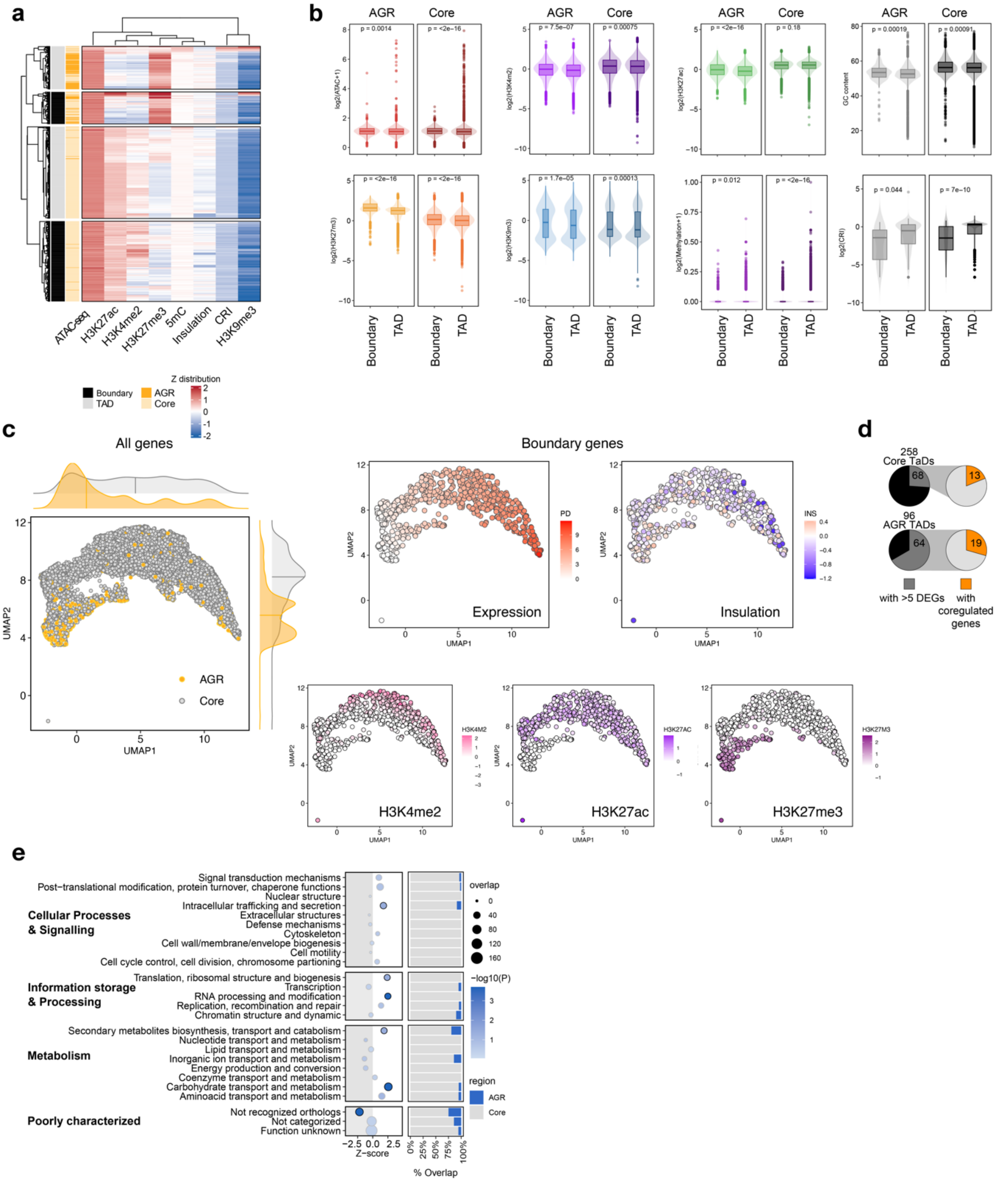


**Figure S3. Chromatin characteristics differentially associate with TADs and boundaries. (a)** k-means clustering ($k$=2) of TADs and boundaries using data presented in Fig. 1E combined with insulation scores and composite repeat-induced point mutation (RIP) index (CRI) values averaged for 100 bp windows and summarized per TAD and boundary region. **(b)** TADs and TAD boundaries differ in histone modifications, CRI, and RIP. P-value after one-way Wilcoxon rank sum test. **(c)** Uniform Manifold Approximation and Projection for Dimensional Reduction (UMAP) to separate genes of *Verticillium dahliae* strain JR2 according to their epigenetic profile, i.e., DNA methylation, CRI (Composite Repeat Index), H3K27ac, H3K27me3, H3K4me2, and H3K9me3. The left top plot displays all genes in the core genome (grey) and in AGRs (yellow). Distributions are shown on top and right of the plot. The remaining plots display only genes located at TAD boundaries and are coloured according to their transcription in PDB (TPM PDB), the insulation score of the boundary they locate in, their H3K4me2 ChIP coverage signal, their H3K4ac ChIP coverage signal, and their H3K27me3 ChIP coverage signal. **(d)** Pie chart showing the proportion of TADs in the core genome and in AGRs containing more than five differentially expressed genes between cultivation for 6 days in PDB or in CZA (grey), and for which DEGs display directionality of differential expression (orange). (**e**) Cluster for orthologs (COGs) gene enrichment in TAD boundaries. The y-axis depicts the COG categories and the x-axis the z-score after a permutation test (10,000 iterations); negative z score indicates depletion, while positive z score shows enrichment. The significance is shown as –log10(FDR adjusted p-value) and is color-coded; circle size is relative to the number of genes overlapping in boundaries per each category. The bar charts on the right indicates the proportion of genes in boundaries that occur in the core genome and AGRs.

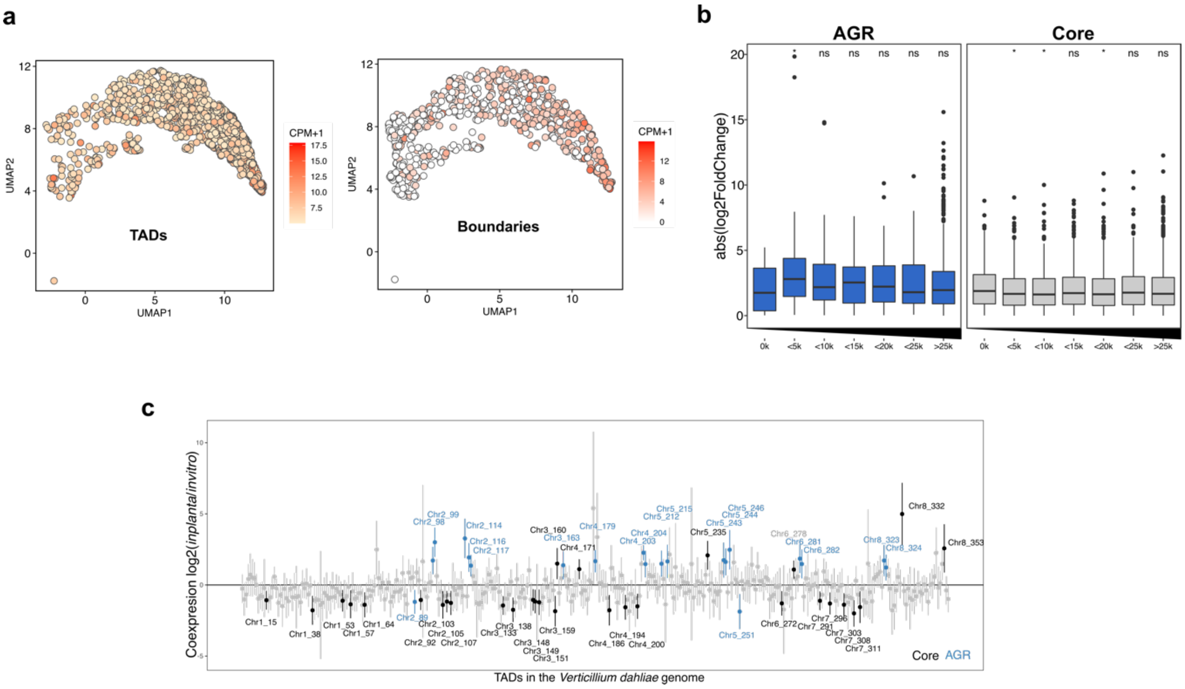

**Figure S4. *In planta* gene expression at TADs and boundaries. (a)** Uniform Manifold Approximation and Projection for Dimensional Reduction (UMAP) to separate genes of *Verticillium dahliae* strain JR2 according to their epigenetic profile and colored based on the gene expression of *V. dahliae* 28 days post inoculation of *Arabidopsis thaliana*. On the left, dots depict genes co-located in TADs, while right plot shows genes co-located in TAD boundaries. Circles are coloured according to the gene expression at 28 dpi of *Arabidopsis thaliana* (TPM AR). (**b**) Absolute log2-fold change in expression between cultivation in PDB and *in planta* (AR) for all genes grouped based on their distance to the closest boundary in the core genome (grey) or in AGRs (blue). Statistically significant differences in average transcription level for the distance groups was compared to the group of genes located in boundaries (distance 0) and determined by the Wilcoxon Rank-Sum test (* $p < 0.05$). Black lines in the boxes depict the median, boxes extend from first to third quartile, vertical lines indicate the 1.5x interquartile range and dots depict outliers for each category. (c) Linear regression effect size of each TAD on differential gene expression between cultivation *in vitro* for 6 days in PDB or *in planta*. Mean effect size of each TAD is shown as a point, with 95% confidence interval, and TADs with a significant effect (95% confidence interval is significantly different from 0) are shown in colour and labelled by corresponding chromosome and TAD number, for TADs in the core genome (black labels) and in AGRs (blue labels).

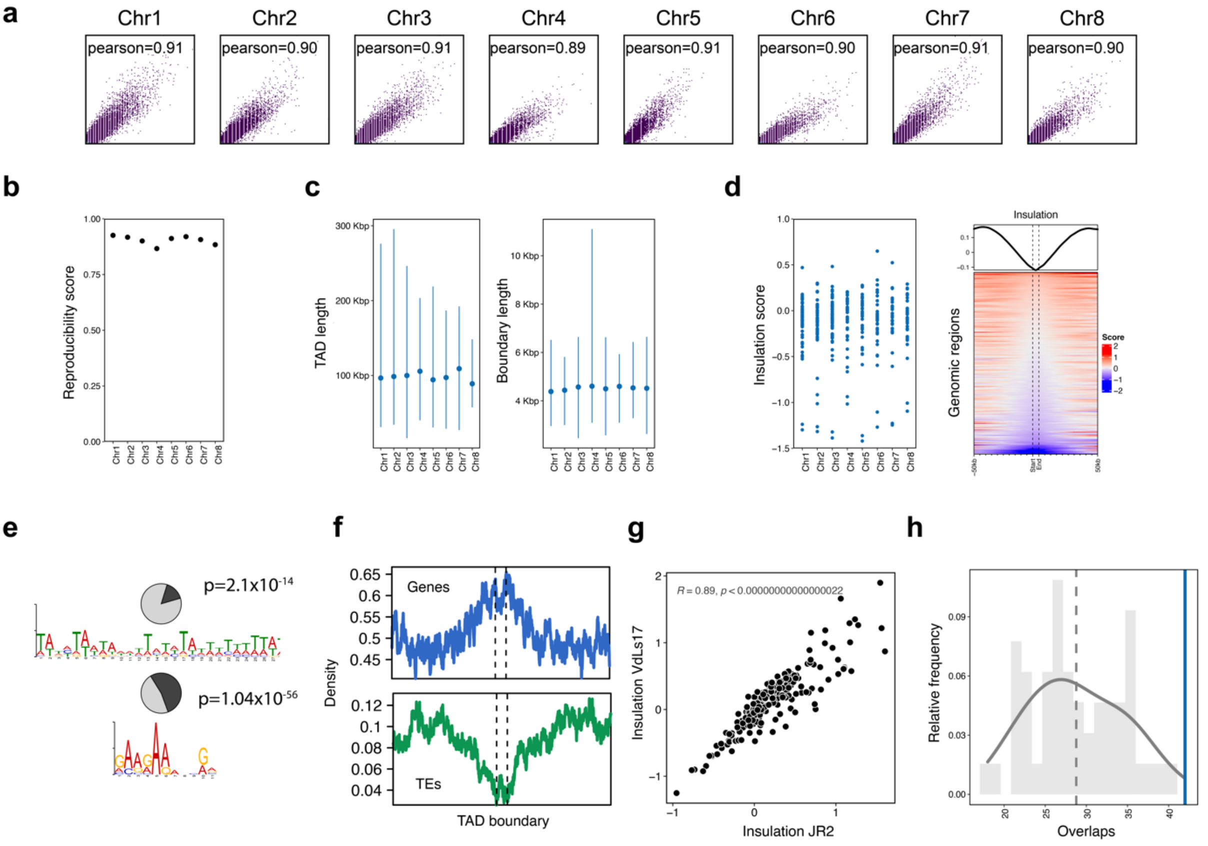


**Figure S5. Characteristics of TADs and boundaries in *Verticillium dahliae* strain VdLs17. (a)** Pearson correlation between replicates of Hi-C interactions for *V. dahliae* strain VdLs17, divided per chromosome. **(b)** Reproducibility score based on a stratified cross-correlation of interactions between replicates of Hi-C interactions for *V. dahliae* strain VdLs17 divided per chromosome. **(c)** Length of TADs and boundaries predicted on the merged replicates, divided per chromosome. **(d)** On the left: Insulation score of boundaries predicted in *V. dahliae* strain VdLs17. On the right: insulation score over boundaries with 50 kb up- and down-stream sequence. **(e)** Presence of the TATA- and GAAG-motifs in boundaries of *V. dahliae* strain VdLs17. **(f)** Enrichment of genes and depletion of transposable elements (TEs) at boundaries in *V. dahliae* strain VdLs17. **(g)** Correlation of insulation score for boundaries predicted in genomic regions that are syntenic between strains JR2 and VdLs17. **(h)** Lowly insulated boundaries of *V. dahliae* strain VdLs17 are enriched in genomic regions that are non-syntenic between JR2 and VdLs17.

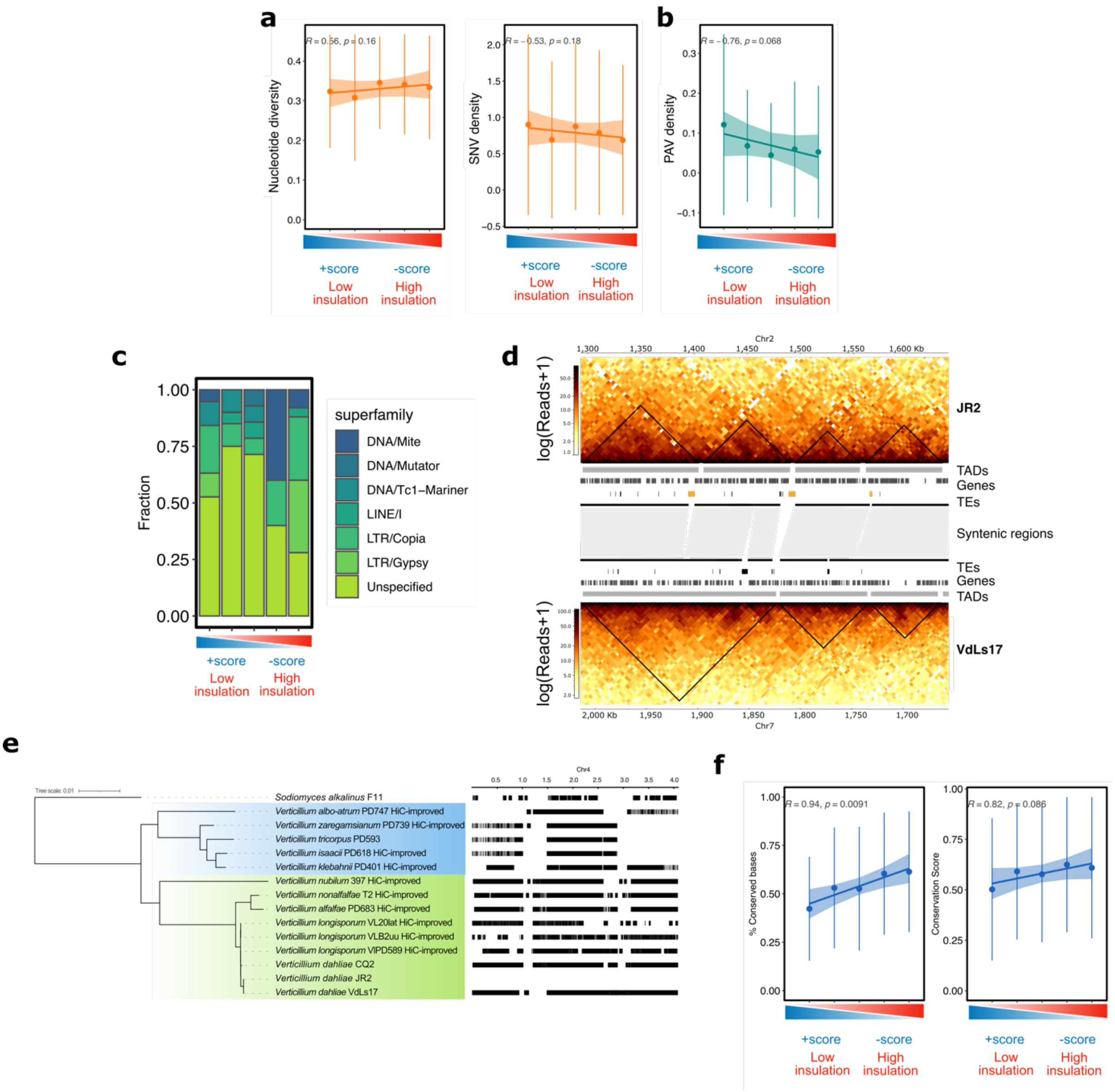


**Figure S6. Conservation of TADs and boundaries in a collection of 42 *Verticillium dahliae* strains and in the *Verticillium* genus. (a)** Correlation between insulation score and nucleotide diversity (left) and SNV density (right) for boundary quintiles separated on their insulation strength. **(b)** Correlation between insulation score and presence/absence variation (PAV) for boundary quintiles separated on their insulation strength. **(c)** Abundance of polymorphic transposable elements (*n*=86) for boundary quintiles separated by their insulation strength in the *V. dahliae* strain JR2. **(d)** Multiple TE insertions in *Verticillium dahliae* strain JR2 coincide with boundary rearrangements between *V. dahliae* strains. Heatmaps represent contact matrixes with TADs (black triangles) over a section of chromosome 2 of JR2 (top) and chromosome 7 of VdLs17 (bottom) that are syntenic. The region occurs inverted in VdLs17 relative to JR2. TADs, genes and TEs are displayed between heatmaps. Synteny between JR2 and VdLs17 indicated as grey blocks. TE insertions in strain JR2 are indicated in yellow. **(e)** Phylogeny of the ten *Verticillium* species using *Sodiomyces alkalinus* to root the phylogeny; Flavexudans and Flavnonexudans clades are depicted in blue and green, respectively. On the left: syntenic regions (black blocks) between *Verticillium* genomes when compared with *V. dahliae* strain JR2, based on chromosome 4 of *V. dahliae* JR2 strain as example. **(f)** Correlation between insulation score and fraction of conserved bases (left) and conservation score (right) for boundary quintiles separated on their insulation strength.

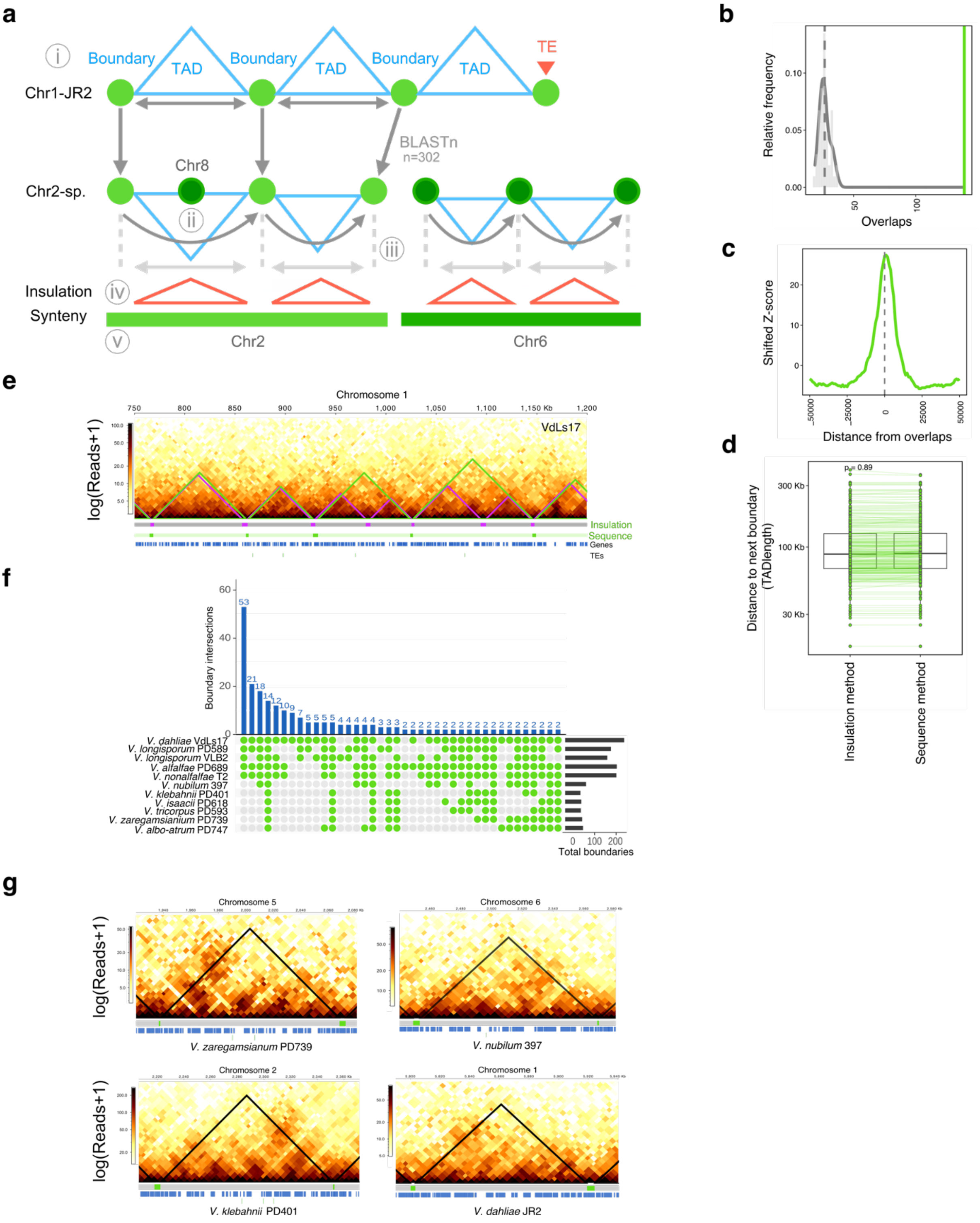


**Figure S7. Predicted TAD boundaries in the *Verticillium* genus display low insulation when compared with adjacent genomic regions. (a)** Overview of our approach to predict the TAD organization in *Verticillium* species with a schematic representation of the *V. dahliae* JR2 reference TAD organization on top, boundaries as green circles and TADs as blue triangles. i) 302 DNA sequences from boundaries (excluding those with TE insertions) were obtained and compared to the genomes of *Verticillium* spp. ii) Contiguous sequences belonging to same chromosome were retained and single interrupting matches from other chromosomes were removed. iii) The distances between contiguous boundaries were calculated in reference and subject. iv) Predicted boundaries were compared to the calculated insulation score from the Hi-C data. v) Predicted boundary locations were compared to the progressive Cactus syntenic regions to crosscheck boundary distribution. **(b)** Distribution of 10,000 iterations of the permutation test for overlaps between the boundaries as predicted for *V. dahliae* strain VdLs17 by the sequence-based method and by the insulation method (grey distribution), the green line indicates significant overlaps between predicted and 'reference' boundaries from *V. dahliae* VdLs17 (Fig. 3), Z-score=27.1264, p=9.9x10-5. **(c)** Z-score shifts from boundaries indicate a high enrichment of overlapping TAD boundaries for the two methods. **(d)** TAD length prediction distribution by the insulation method and the sequence method is not significantly different. P-value after one-way Wilcoxon rank sum test. **(e)** Overlap in TAD prediction for strain VdLs17 using the insulation method (purple triangles) and the JR2-sequence-based method (green triangles), with partial chromosome 1 of the strain VdLs17 (755,315-1167941 bp) as example. Tracks depict in grey and purple, TADs and boundaries by the insulation method respectively; in light green and green, TADs and boundaries by the sequence-based method respectively; Genes in blue and TEs in green. **(f)** TAD boundaries predicted in the ten *Verticillium* species ordered according to JR2-sequence-based method, on the bottom-right the total number of boundaries predicted in each species. On the bottom, a combination matrix depicting combinations of species (in green) in which boundaries are shared. On the top, the total amount of boundaries shared for each species combination. **(g)** TADs are conserved in syntenic regions in the *Verticillium* genus. One syntenic TAD is shown in four species of the *Verticillium* genus. Black tringle indicates the TAD. Grey and green tracks depict TADs and boundaries respectively. Genes in blue and TEs in green.

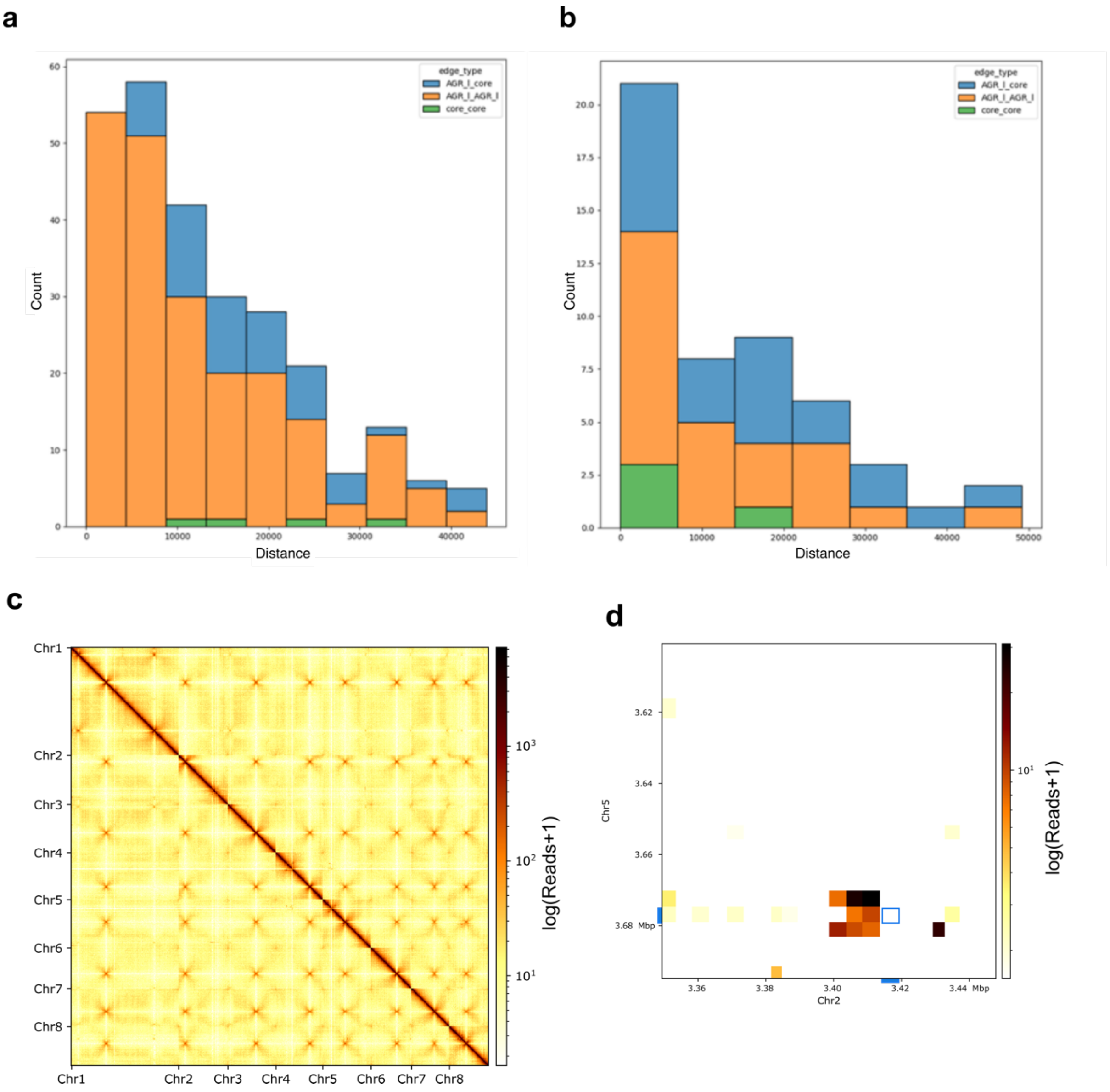


**Figure S8. Distance between colocalizing regions and neighboring duplicated regions in *Verticillium dahliae*. (a)** Duplicated regions occur in close proximity to physical co-located regions in *V. dahliae* JR2. Y-axis depict distance from co-located regions, and colors show the type of physical co-location event (AGR-to-Core, AGR-to-AGR, Core-to-Core). **(b)** Duplicated regions occur near physically co-located regions in *V. dahliae* VdLs17. Y-axis depict distance from co-located regions, and colors show the type of physical co-location event (AGR-to-core, AGR-to-AGR, core-to-core). Colocalization events involving AGR-to-core regions are shown in blue, AGR-to-AGR regions in orange and core-to-core regions in green. (**c**) Verticillium dahliae JR2 Hi-C matrix showing the high interaction within chromosomes and between centromeric regions, suggesting a Rabl configuration. (d) High interaction between AGRs in chromosome 5 and 2 occurs in close proximity to segmental duplications (blue box).

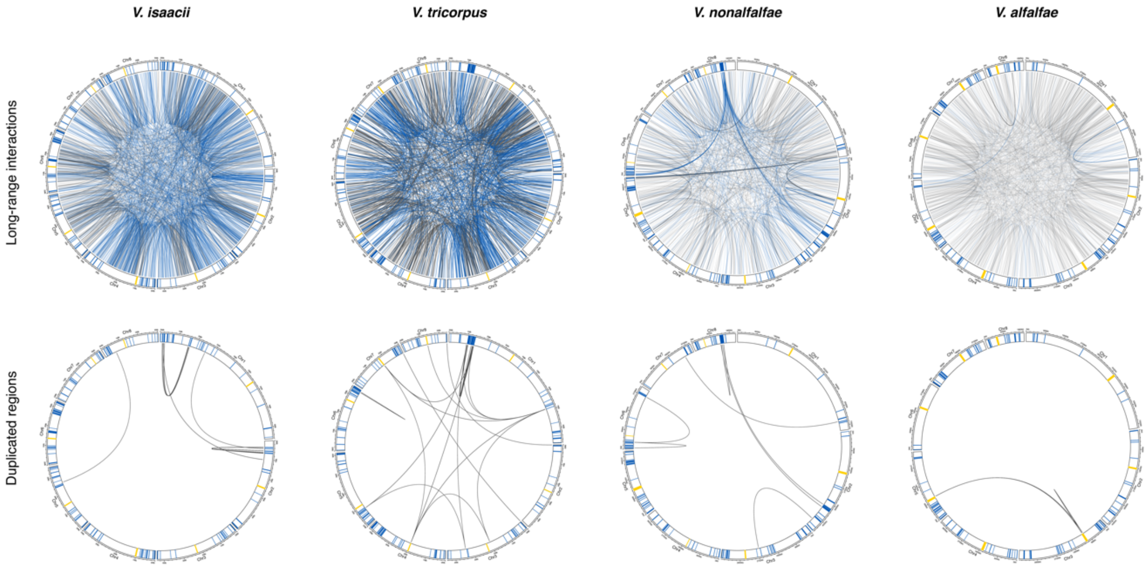


**Figure S9. Non-centromeric long-range interactions co-locate with duplicated regions in the *Verticillium* genus.** All circular plots display the chromosomes with centromeres in yellow, AGR regions in blue, and core regions in white. For every genome, the upper plot shows non-centromeric long-range interactions that exceed the average interaction strength of centromeres as edges and are color-coded according to the respective genomic annotation of the regions they connect, with AGR interactions in blue and core interactions in grey. In the lower plot, the edges represent segmental duplications.

**Table S1. Aggregated counts of long-range interactions in *Verticillium dahliae* and their overlap with segmental duplications**

| | *V. dahliae* strain JR2 | *V. dahliae* strain VdLs17 |
|---|---|---|
| Genome-wide | 1842 | 1978 |
| Centromere | 953 | 527 |
| Non-centromere | 889 | 1451 |
| AGR | 475 | 452 |
| Core-AGR | 225 | 407 |
| AGR-AGR | 250 | 45 |
| Core-core | 414 | 999 |
| Segmental duplication | 264 | 50 |
| AGR-AGR | 207 | 21 |
| Core-AGR | 53 | 20 |

**Table S2. Expected and observed non-centromeric colocalization events between genetically distant regions in *Verticillium*.**

| Species | Non-centromeric colocalization events | Expected core | Observed core | Expected AGR | Observed AGR | p-value[a] |
|---|---|---|---|---|---|---|
| *V. dahliae* JR2 | 889 | 880 | 639 | 162 | 475 | 6.8695E-148 |
| *V. dahliae* VdLs17 | 1451 | 1442 | 999 | 217 | 452 | 6.1638E-87 |
| *V. albo-atrum* | 897 | 893 | 880 | 111 | 143 | 0.002152792 |
| *V. klebahnii* | 573 | 571 | 571 | 65 | 178 | 1.24565E-44 |
| *V. nubilum* | 582 | 578 | 572 | 81 | 173 | 1.52804E-24 |
| *V. nonalfalfae* | 891 | 887 | 873 | 105 | 157 | 3.46163E-07 |
| *V. isaacii* | 1645 | 1639 | 1632 | 191 | 132 | 1.93215E-05 |
| *V. alfalfae* | 1455 | 1450 | 1453 | 163 | 71 | 5.74392E-13 |
| *V. tricorpus* | 1053 | 1048 | 1051 | 126 | 73 | 2.32935E-06 |
| *V. longisporum* | 2426 | 2420 | 2407 | 231 | 234 | 0.741519345 |
| *V. longisporum* | 1201 | 1198 | 1195 | 110 | 91 | 0.069731441 |

[a]Based on chi-square test between observed and expected colocalization events in the different genomic compartments.

**Table S3. Observed and expected association between genomic colocalization events among genetically distant regions and duplicated genomic regions.**

| Species | Observed duplication-colocalizations | Expected duplication-colocalizations[a] | Duplication-colocalizations core | Duplication-colocalizations AGR | p-value[b] |
|---|---|---|---|---|---|
| *V. dahliae* JR2 | 264 | 0.38 | 57 | 260 | 1.06E-267 |
| *V. dahliae* VdLs17 | 50 | 0.54 | 29 | 41 | 4.21E-184 |
| *V. albo-atrum* | 11 | 0.23 | 8 | 6 | 2.93E-133 |
| *V. klebahnii* | 24 | 0.06 | 24 | 0 | 3.60E-200 |
| *V. nubilum* | 16 | 0.69 | 15 | 13 | 6.92E-132 |
| *V. nonalfalfae* | 6 | 0.07 | 3 | 5 | 4.01E-137 |
| *V. isaacii* | 13 | 0.11 | 7 | 6 | 1.03E-157 |
| *V. alfalfae* | 0 | 0 | 0 | 0 | 1.00E+00 |
| *V. tricorpus* | 4 | 0.09 | 4 | 2 | 2.35E-114 |
| *V. longisporum* | 2400 | 3.63 | 2381 | 232 | 0.00E+00 |
| *V. longisporum* | 296 | 1.82 | 295 | 9 | 3.19E-235 |

[a]Average of 100 permutations.
[b]Based on t-test of the observed duplicated regions neighboring colocalization events versus the expected distribution (100 permutations).

**Table S4. Association between genomic colocalization events among genetically distant regions and duplicated genomic regions.**

| Species | AGR-AGR | AGR-core | core-core | Total |
|---|---|---|---|---|
| *V. dahliae* JR2 | 207 | 53 | 4 | 264 |
| *V. dahliae* VdLs17 | 21 | 20 | 9 | 50 |
| *V. albo-atrum* | 3 | 3 | 5 | 11 |
| *V. klebahnii* | 0 | 0 | 24 | 24 |
| *V. nubilum* | 1 | 12 | 3 | 16 |
| *V. nonalfalfae* | 3 | 2 | 1 | 6 |
| *V. isaacii* | 6 | 0 | 7 | 13 |
| *V. alfalfae* | 0 | 0 | 0 | 0 |
| *V. tricorpus* | 0 | 2 | 2 | 4 |
| *V.longisporum* | 19 | 213 | 2168 | 2400 |
| *V. longisporum* | 1 | 8 | 287 | 296 |
